# Self-generated whisker movements drive state-dependent sensory input to developing barrel cortex

**DOI:** 10.1101/2020.01.21.914259

**Authors:** James C. Dooley, Ryan M. Glanz, Greta Sokoloff, Mark S. Blumberg

## Abstract

Cortical development is an activity-dependent process [1–3]. Regarding the role of activity in developing somatosensory cortex, one persistent debate concerns the importance of sensory feedback from self-generated movements. Specifically, recent studies claim that cortical activity is generated intrinsically, independent of movement [3, 4]. However, other studies claim that behavioral state moderates the relationship between movement and cortical activity [5–7]. Thus, perhaps inattention to behavioral state leads to failures to detect movement-driven activity [8]. Here, we resolve this issue by associating local field activity (i.e., spindle bursts) and unit activity in the barrel cortex of 5-day-old rats with whisker movements during wake and myoclonic twitches of the whiskers during active (REM) sleep. Barrel activity increased significantly within 500 ms of whisker movements, especially after twitches. Also, higher-amplitude movements were more likely to trigger barrel activity; when we controlled for movement amplitude, barrel activity was again greater after a twitch than a wake movement. We then inverted the analysis to assess the likelihood that increases in barrel activity were preceded within 500 ms by whisker movements: At least 55% of barrel activity was attributable to sensory feedback from whisker movements. Finally, when periods with and without movement were compared, 70–75% of barrel activity was movement-related. These results confirm the importance of sensory feedback from movements in driving activity in sensorimotor cortex and underscore the necessity of monitoring sleep-wake states to ensure accurate assessments of the contributions of the sensory periphery to activity in developing somatosensory cortex.

## Results

Previous studies have identified behavioral state as a factor that moderates the relationship between self-generated movements and neural activity. Specifically, in infant rats before postnatal day (P) 11, whereas sensory feedback from limb movements during wake only weakly activates sensorimotor cortex, sensory feedback from limb twitches during active sleep triggers strong cortical responses [5–7]. However, in many studies of activity in developing somatosensory cortex, either behavioral state was not measured [4, 9, 10] or the subjects were anesthetized [11–13]. It is plausible, then, that by ignoring behavioral state, we underestimate the importance of sensory feedback from self-generated movements for driving early neural activity.

To address this issue directly, we recorded whisker movements during active sleep (AS) and wake (W) in P5 rats while recording extracellular activity in the C row of barrel cortex. All whiskers, except those in the C row, were trimmed. Whisker movements were recorded using high-speed video at 100 frames•s^−1^ and analyzed using DeepLabCut, a deep-learning method for markerless tracking of behavior (Figure 1A; [14, 15]). Local field potential (LFP) and spike trains of individual neurons were extracted from the raw neural data (Figure 1B-D). LFP recordings were filtered for spindle bursts (8–40 Hz), the predominant oscillation observed in developing sensorimotor cortex [1, 9].

**Figure 1.**
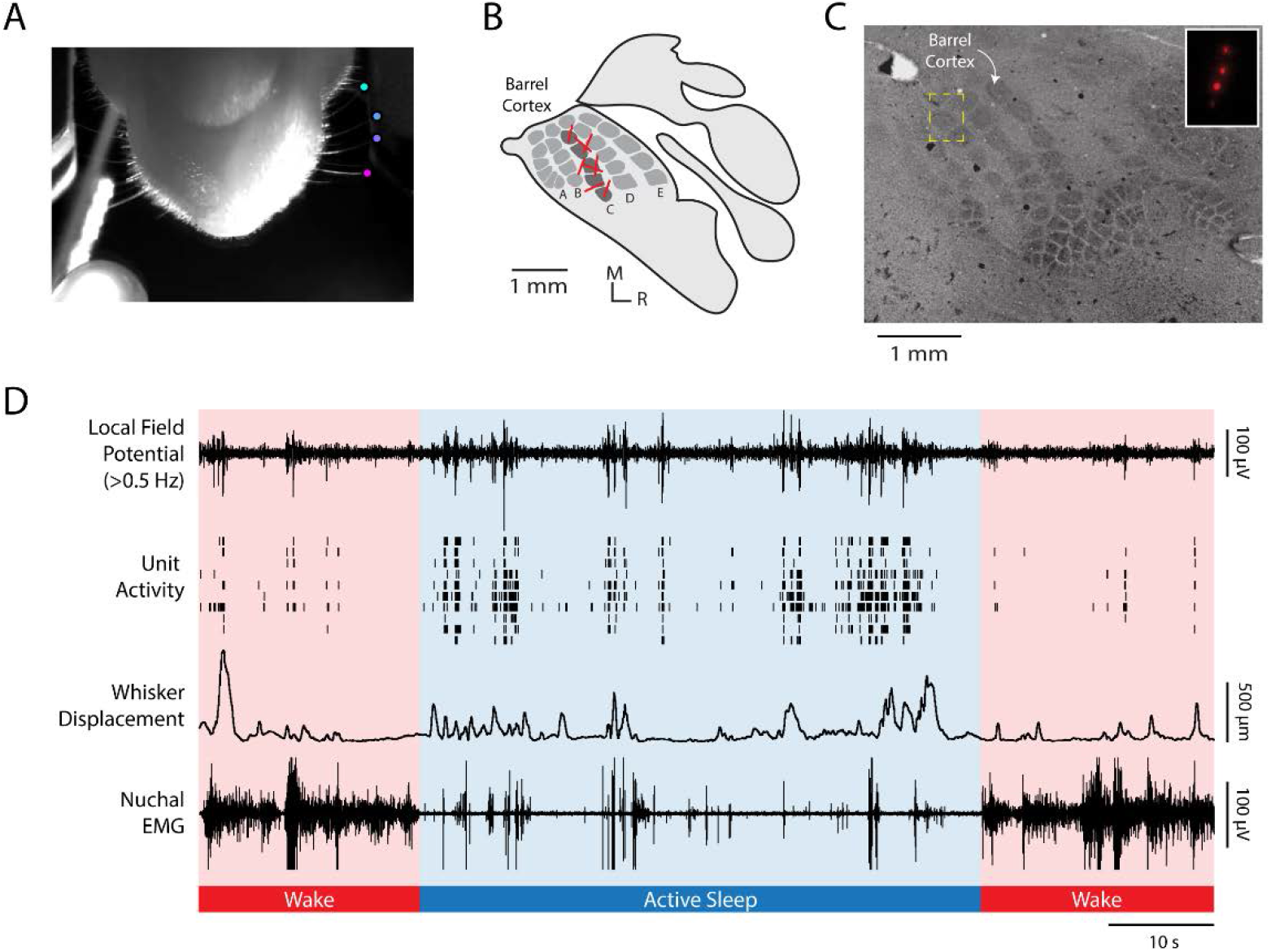
Methods and representative recording (A) video frame, shot from below, illustrating the method for tracking whisker movements using DeepLabCut. Colored dots are located at the tips of four C-row whiskers. Whisker displacement was measured along the animal’s anterior-posterior axis. At the bottom left of the image, the LED used for data synchronization is shown. (B) Illustration of primary somatosensory cortex (S1) in a P5 rat. Whisker barrels are depicted in dark gray within the S1 map. Electrode shank recording sites for each animal are represented by red lines in the C row of the barrel field. (C) Representative histological section of barrel cortex. The S1 barrel field was visualized using cytochrome oxidase. The electrode location for a single pup is shown within the yellow dashed box, which is enlarged in the inset to show the fluorescent electrode tracts. (D) Representative 100-s record for an individual pup during active sleep (blue shading) and wake (red shading). From top to bottom: *Local field potential* (LFP) recorded in barrel cortex. *Unit activity* recorded from barrel cortex; each row denotes a different single unit. *Whisker displacement*, in μm, of all tracked C-row whiskers (i.e., mean displacement). *Nuchal EMG* recording indicates periods of high muscle tone indicative of wake and periods of low muscle tone, punctuated by brief spikes (i.e., nuchal twitches), indicative of active sleep. Note that LFP and unit activity are highest during active sleep and that whisker movements coincide with increases in barrel cortex activity.

Animals slept for 32.7 ± 1.93 minutes during the 60-min recording sessions (Figure S1A). Whisker twitches and wake movements occurred at equal rates (Figure S1B; AS: 27.1 ± 1.45 min^−1^; W: 25.6 ± 2.28 min^−1^; t(7) = 0.43, p = .679, adj. 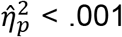). Whisker twitches and wake movements also had similar displacements and durations (Figure S1C).

If whisker movements and barrel cortex activity are causally unrelated, we would expect them to sometimes occur together by chance. Therefore, the analyses below compare the relationship between whisker movements and barrel cortex activity with the expected relationship due to chance.

### Whisker movements drive neural activity

We first measured the change in LFP power (2–100 Hz) before and after whisker twitches and wake movements. As shown in Figure 2A, peak LFP power increased significantly in the 500-ms period after whisker movements. Peak power was greater after twitches (7.8 ± 0.62 dB) than wake movements (5.8 ± 0.58 dB; t(7) = 4.19, p = .004, adj. 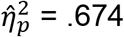). In addition, as shown in Figure 2B, the rate of spindle bursts was significantly higher during active sleep (7.9 ± 0.75 min^−1^) than during wake (3.3 ± 0.30 min^−1^; t(7) = 5.95, p = .001, adj. 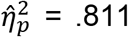). The mean amplitude, duration, and peak frequency of spindle bursts produced during active sleep and wake did not differ significantly (data not shown).

**Figure 2.**
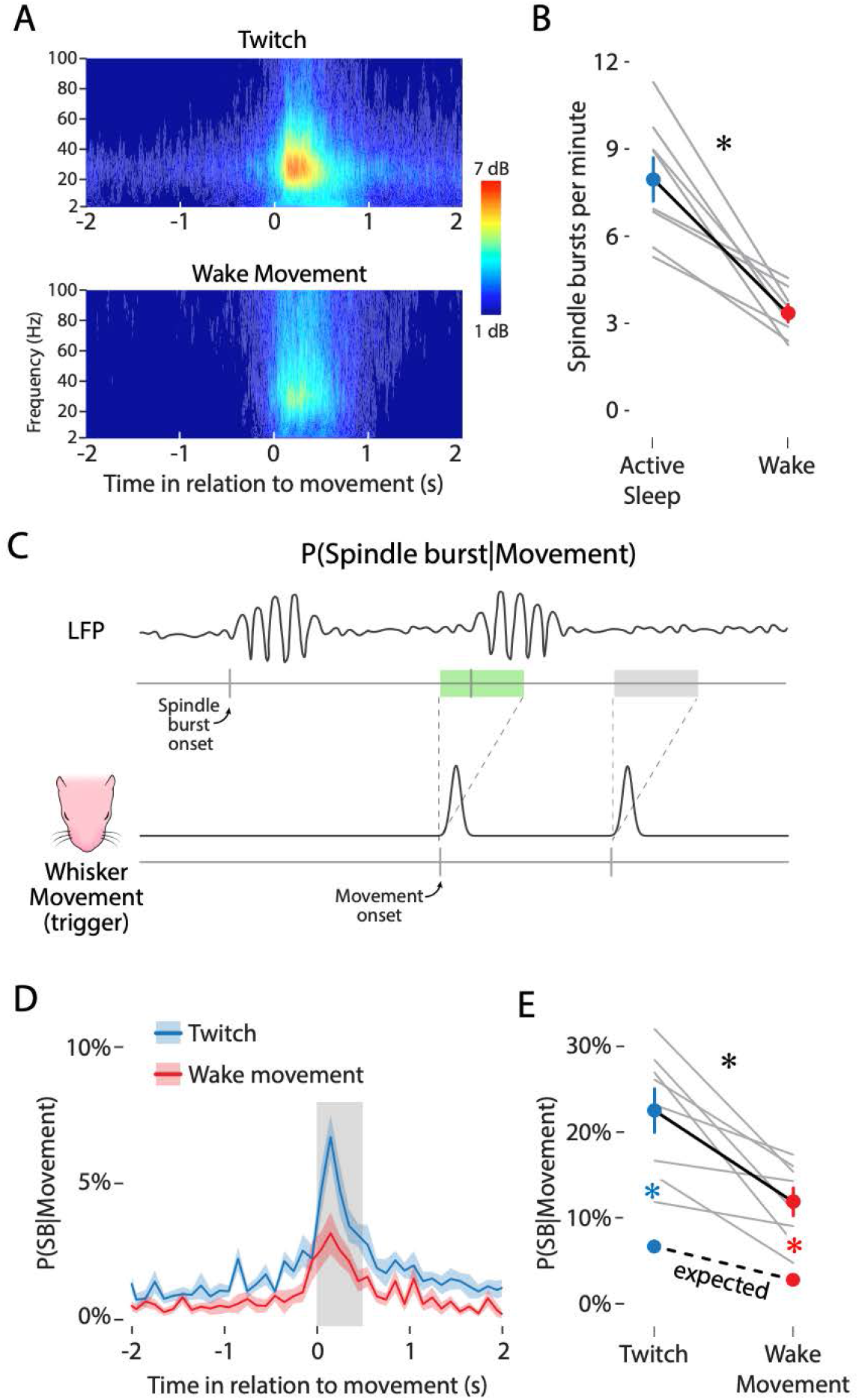
Whisker movements drive spindle bursts in barrel cortex (A) Frequency spectrograms showing LFP activity, averaged across pups, in relation to the onset (at 0 s) of whisker twitches (top) and wake movements (bottom). The color axis represents LFP power, in dB above baseline. (B) Mean (±SEM) rate of spindle bursts per minute during active sleep (blue circle) and wake (red circle). Gray lines show spindle burst rate for individual pups (N = 8). Asterisk denotes significant difference, p < .05. (C) Illustration of the method used to determine the likelihood of a spindle burst given a whisker movement. Bottom: The black line represents the displacement of the whiskers as a function of time and the two gray tick marks below indicate the onset of whisker movements. Each movement onset initiates a 500-ms window, shown above. Top: LFP signal with two spindle bursts. The tick marks along the gray line below indicate the onset of a spindle burst. For this analysis, two key possibilities are shown: a whisker movement that is followed within 500 ms by a spindle burst (green rectangle) and a movement that is not followed within 500 ms by a spindle burst (gray rectangle). (D) Mean (±SEM) likelihood of a spindle burst given a whisker twitch (blue line) or wake movement (red line) in relation to movement onset. The gray rectangle after movement onset shows a 500-ms window. Note that the peak increase in spindle burst likelihood for both twitches and wake movements falls within this 500-ms window. (E) Mean (±SEM) likelihood of a spindle burst within 500 ms of a twitch (upper blue circle) or wake movement (upper red circle). The lower colored circles represent the mean (±SEM) likelihood of a spindle burst due to chance during active sleep (blue) and wake (red). Gray lines show data for individual pups (N = 8). The expected spindle burst likelihoods for individual animals are not shown. The black asterisk indicates a significant interaction (p < .05). Colored asterisks indicate significant observed-expected likelihood differences (p < .05) for whisker twitches (blue) and wake movements (red).

We next measured the likelihood of observing a spindle burst in barrel cortex in the 500-ms period after a whisker movement (Figure 2C). A 500-ms time window was chosen because it captures the period of increased spindle burst likelihood after a whisker movement (Figure 2D). After a whisker twitch, the observed likelihood of a spindle burst was 22.5 ± 2.5%, compared with the expected value of 6.7 ± 0.6% (Figure 2E, blue dots).

In contrast, after a wake movement, the observed likelihood of a spindle burst was 11.9 ± 1.6%, compared with the expected value of 2.8 ± 0.3% (Figure 2E, red dots). Spindle bursts were significantly more likely to be triggered by twitches than wake movements, as evidenced by a significant interaction between behavioral state and the observed-expected likelihood difference (F(1, 7) = 8.62, p = .022, adj. 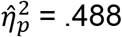). In summary, these results indicate that spindle bursts are twice as frequent during active sleep and 3–4 times more likely to occur after whisker movements than expected by chance.

Parallel analyses performed on single units in barrel cortex yielded similar results. Mean firing rates were higher during active sleep than wake (Figure S2A; t(36) = 6.31, p < .001, adj. 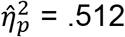). After twitches, firing rates increased to a greater degree than after wake movements (Figure S2B; t(36) = 3.84, p < .001, adj. 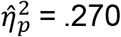). Additionally, the observed-expected likelihood of a firing rate increase was significantly greater after twitches than wake movements (Figure S2C; F(1, 36) = 10.39, p = .003, adj. 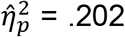). Finally, there was a strong relationship between spindle bursts and unit activity: When twitches or wake movements triggered spindle bursts, we observed substantially stronger single-unit responses than when movements did not trigger spindle bursts (Figure S2D, E).

### Movement amplitude moderates neural activity

That a small minority of movements trigger a response in barrel cortex recently led researchers to conclude that barrel activity is mostly independent of self-generated movements (see [4]). However, this conclusion is confounded by an important statistical issue: At these ages, whisker movements are relatively abundant (>25 min^−1^; Figure S1B) compared with neural activity (3-8 spindle bursts•min^−1^, Figure 2B; <1 spike•s^−1^, Figure S2A). Clearly, many whisker movements do not trigger neural activity. However, not all whisker movements are the same; specifically, small whisker movements may be less likely than large ones to trigger barrel cortex activity.

Thus, we next analyzed whether movement amplitude moderates the likelihood of barrel cortex activity. To test this possibility, whisker movements were assigned to ten bins, from smallest to largest peak displacement. Small movements were much more frequent than large movements (Figure S3A) and the mean amplitude of twitches and wake movements was equal for each bin (Figure S3B).

The likelihood of a spindle burst occurring within the 500-ms period after a whisker twitch or wake movement was determined for each bin (Figure 3A). Spindle burst likelihood in barrel cortex increased with movement amplitude (Figure 3B). The likelihood of a spindle burst occurring after the smallest-amplitude whisker twitches was 3.7 ± 0.4% and increased significantly with amplitude to 48.1 ± 8.3% (Figure 3B, blue line). For wake movements, the likelihood of a spindle burst also increased with amplitude, from 1.4 ± 0.2% to 20.6 ± 2.9% (Figure 3B, red line). This amplitude-dependent increase was greater for twitches than for wake movements, as evidenced by a significant interaction between behavioral state and movement amplitude (F(9, 63) = 2.34, p = .024, adj. 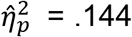). In contrast with movement amplitude, movement duration was not associated with increased spindle burst likelihood (data not shown).

**Figure 3.**
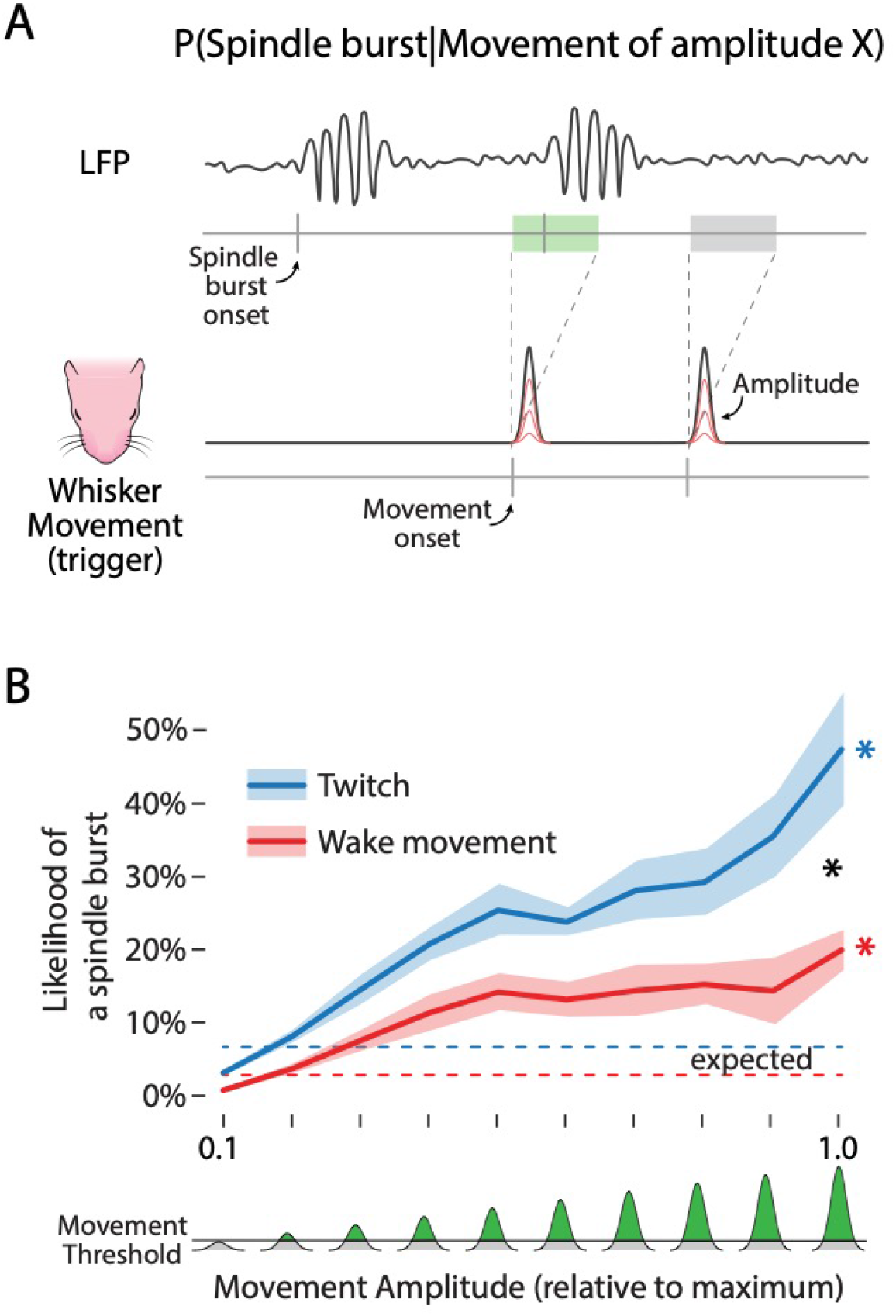
Movement amplitude moderates spindle burst activity (A) Illustration of the method used to determine the likelihood of a spindle burst given a whisker movement of amplitude X. Illustration is identical to that in Figure 2C, except that the red lines indicate whisker displacements of varying amplitude. (B) Mean (±SEM) likelihood of a spindle burst given a twitch (blue line) or wake movement (red line) of a given amplitude. Whisker movements were sorted into one of ten bins of increasing amplitude (0-1 along the x-axis), normalized to the maximum amplitude of whisker deflections observed in each pup. *Bottom:* The amount of whisker displacement, relative to the maximum displacement for each pup, depicted in green. The horizontal line denotes the threshold used to detect whisker movements (see Methods). The expected likelihood of a spindle burst for movements of any amplitude is depicted as a dashed blue (active sleep) or dashed red (wake) line. Black asterisk denotes a significant interaction (p < .05). Colored asterisks indicate significant observed-expected likelihood differences (p < .05) for whisker twitches (blue) and wake movements (red).

We observed a similar relationship between movement amplitude and unit activity in barrel cortex (Figure S3C). The likelihood of unit activity increased with movement amplitude. This increase was stronger for twitches than wake movements, as evidenced by a significant interaction between behavioral state and movement amplitude (F(4.23, 152.39) = 4.67, p < .001, adj. 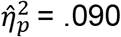).

These results show that although large-amplitude movements represent a small proportion of all movements, they more reliably trigger barrel activity. However, even the largest amplitude movements trigger a spindle burst only 48% of the time during active sleep, and 21% of the time during wake. The probabilistic nature of sensory responses to movement is puzzling as it suggests that cortical responses are modulated both within and between behavioral states. Additional work is needed to characterize the modulatory mechanisms involved and determine their consequences for neural plasticity.

### Neural activity is reliably preceded by whisker movements

As noted above, the likelihood of a whisker movement producing a spindle burst is necessarily low as a result of the high rate of movement and the low rate of neural activity. To more fully assess the relationship between movement and spindle bursts, we inverted the analysis and determined the likelihood that a given spindle burst was preceded by a movement (Figure 4A). By analogy, to understand the relationship between smoking (a relatively high-frequency event) and lung cancer (a relatively low-frequency event), one would assess both the likelihood that smokers will develop lung cancer and the likelihood that people with lung cancer smoked.

**Figure 4.**
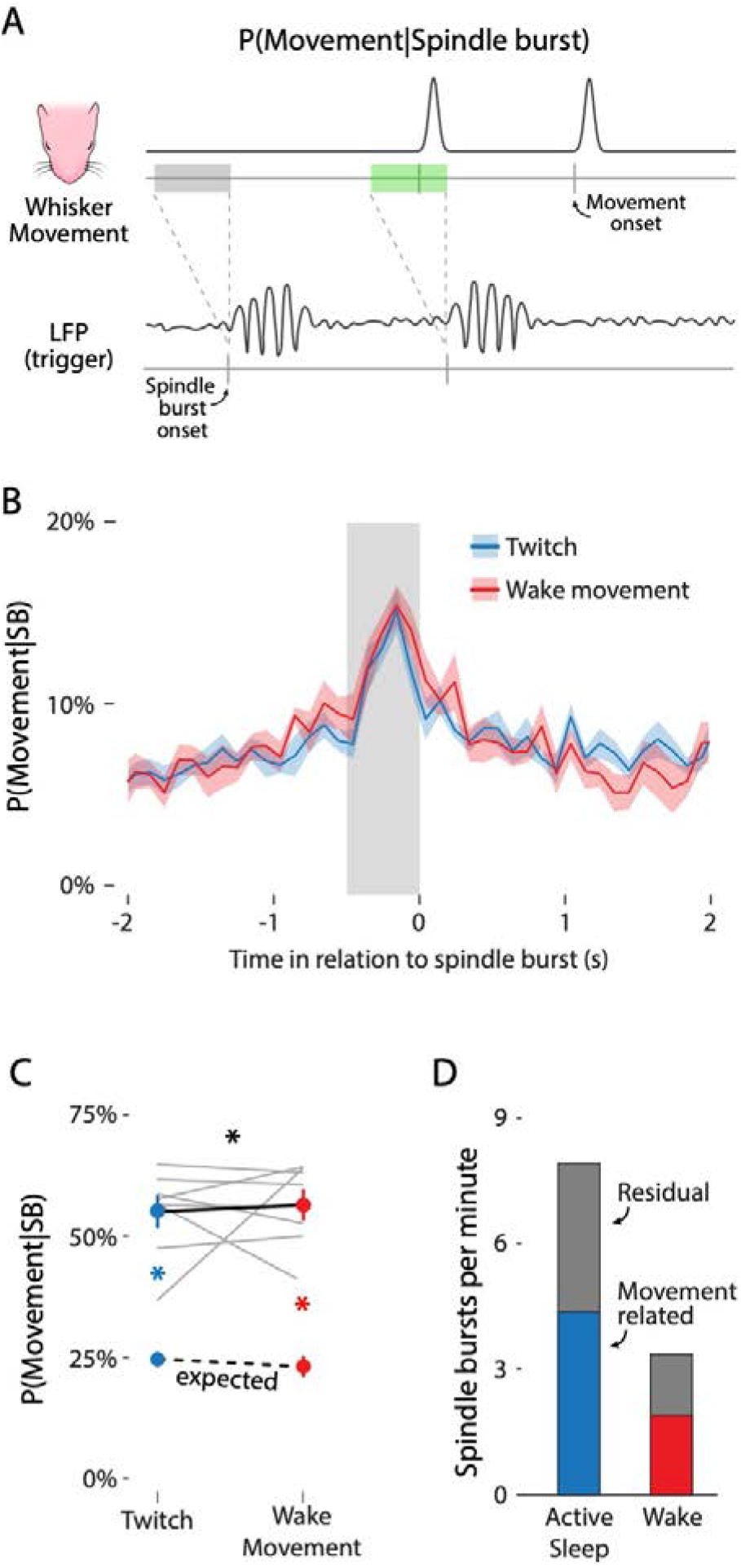
Spindle bursts are reliably preceded by whisker movements (A) Illustration of the method used to determine the likelihood of a whisker movement given a spindle burst. Illustration is the reverse of that in Figure 2C, with the triggering event being the onset of a spindle burst (bottom) rather than a whisker movement. For this analysis, two key possibilities are shown: a spindle burst that is preceded within 500 ms by a whisker movement (green rectangle) and a spindle burst that is not preceded within 500 ms by a whisker movement (gray rectangle). (B) Mean (±SEM) likelihood of a whisker twitch (blue) or wake movement (red) in relation to the onset of a spindle burst. The gray rectangle before spindle burst onset shows a 500-ms window. (C) Mean (±SEM) likelihood of a whisker twitch (upper blue circle) or wake movement (upper red circle) within the 500-ms window before a spindle burst. The lower colored circles represent the mean (±SEM) likelihood of a twitch (blue) or wake movement (red) due to chance. Gray lines show data for individual pups (N = 8). The expected movement likelihood for individual pups is not shown. Black asterisk indicates a significant interaction (p < .05). Colored asterisks indicate significant observed-expected likelihood differences (p < .05) for whisker twitches (blue) and wake movements (red). (D) Mean rate of spindle bursts per minute during active sleep (blue bar) and wake (red bar) directly attributable to whisker movements (i.e., *movement-related* activity). Gray bars represent the rate of spindle bursts not attributable to whisker movement (i.e., *residual* activity).

As with the previous analyses, we observed the highest likelihood of a whisker movement within a 500-ms window before spindle burst initiation during both active sleep and wake (Figure 4B). The likelihood of a whisker movement preceding a spindle burst was significantly greater than the expected value due to chance (Figure 4C; F(1, 7) = 163.45, p < .001, adj. 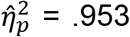). Whisker twitches preceded 55.0 ± 3.1% of all spindle bursts during active sleep, compared with the expected value of 25.0 ± 1.3%. Wake movements preceded 56.4 ± 2.9% of all spindle bursts during wake, compared with the expected value of 23.2 ± 2.0%. There was no effect of behavioral state: Spindle bursts during sleep or wake were equally likely to be preceded by a twitch or wake movement, respectively. However, when expressed in raw numbers (Figure 4D), more than twice as many spindle bursts were preceded by twitches than wake movements.

We again observed similar results for unit activity (Figure S4). Proportionately, the percentage of unit activity preceded by twitches (53.8 ± 2.6%) was lower than the percentage of activity preceded by wake movements (60.4 ± 3.4%), indicated by a significant interaction between behavioral state and the observed-expected amount (F(1,7) = 12.12, p = .010, adj. 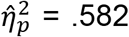). Nevertheless, significantly more unit activity than expected was preceded by both twitches (t(7) = 13.70, p < .001, adj. 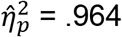) and wake movements (t(7) = 13.17, p < .001, adj. 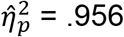).

### Neural activity increases during periods of movement

The 45% of residual activity identified in Figure 4D could arise if sensory feedback from movements in other parts of the body were able to drive activity in barrel cortex. Alternatively, the residual activity could arise spontaneously within the brain, independent of movement. To address these two possibilities, we used a time-based behavioral analysis that includes movements of the ipsilateral whiskers, hindlimbs, and tail. Specifically, we classified each timepoint into one of three movement categories based on whether (1) the contralateral whiskers were moving, (2) the contralateral whiskers were not moving but other body parts were moving, and (3) the pup was not detectably moving. Each period of movement began with the onset of the first movement and ended 500 ms after the last movement. As shown in Figure 5A, pups spent roughly half their time moving or not moving, and during periods when pups were moving, the contralateral whiskers were involved the majority of the time.

**Figure 5.**
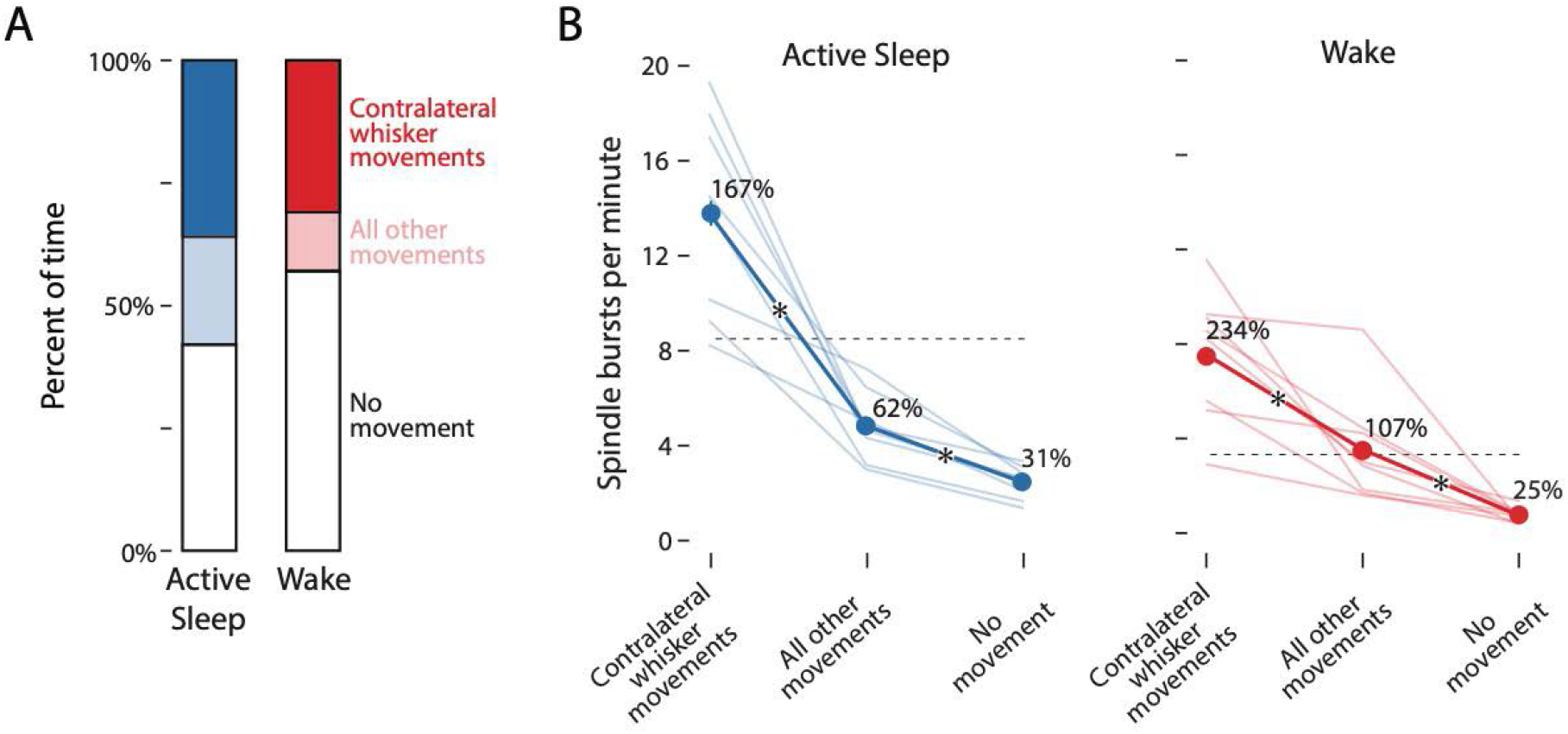
Neural activity increases during periods of movement (A) Mean percent of time during active sleep and wake when the contralateral whiskers were moving (dark blue/red), when other parts of the body were moving but the contralateral whiskers were not (light blue/red), and when no discernible movements were detected (white). (B) Mean (±SEM) rate of spindle bursts per minute during active sleep (left, blue) and wake (right, red). Light blue/red lines show data for individual pups (N = 8). Dashed horizontal lines indicate the mean rate of spindle bursts across the entirety of active sleep and wake periods. Adjacent to each mean value, spindle burst rate is expressed as a percent change in relation to the mean rate within that state. Asterisks indicate significant differences (p < .05) between movement categories.

As shown in Figure 5B, the rate of spindle burst production was influenced both by movement category (F(2, 14) = 6.33, p < .001, adj. 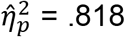) and behavioral state (F(1, 7) = 6.33, p = .040, adj. 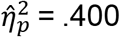). For both active sleep and wake, periods of contralateral whisker movements were associated with the highest rates of spindle burst production (AS: 13.8 ± 1.48 min^−1^; W: 7.5 ± 0.98 min^−1^). Periods of other movements corresponded to an intermediate rate of spindle burst production (AS: 4.8 ± 0.51 min^−1^; W: 3.5 ± 0.82 min^−1^). Periods of no detectable movement were associated with the lowest rate of spindle burst production (AS: 2.5 ± 0.24; W: 0.8 ± 0.11 min^−1^). Because periods of other movements elevated the rate of spindle burst production above the level seen in the no movement periods, these results suggest some imprecision in the somatotopic organization of barrel cortex at this age, as described previously [17].

Finally, when expressed as a percentage of the overall spindle burst rate (Figure 5B, dotted lines), the rate of spindle bursts during periods of no movement was 30.6 ± 2.6% during active sleep and 24.8 ± 4.0% during wake. Accordingly, self-generated movements account for approximately 70% of spindle bursts during active sleep and 75% during wake. These values represent our current best estimate of the percentage of spindle bursts in barrel cortex that are related to movement.

## Discussion

Before the discovery of retinal waves, researchers debated whether visual system development is “predetermined” (e.g., see [16–18]). Today, this issue is largely settled: There is an overwhelming consensus that the input provided by retinal waves is critical to typical development of the visual system [19]. Indeed, rather than debating whether retinal inputs shape the visual system (i.e., whether these inputs are permissive), the field has moved on to asking how patterns of retinal activity lead to normal retinotopy across all levels of the visual system (i.e., how these inputs are instructive; see [20]).

In the somatosensory system, however, the debate continues about the importance of peripheral sensory input for cortical development. Whereas some groups posit that self-generated movements, particularly twitches, drive critical early activity in somatosensory cortex [5, 9], others explicitly state that the sensory feedback from movements is not necessary, instead positing that spontaneous neural activity within the brain is alone sufficient for typical development [3, 4].

It is difficult to imagine how spontaneous neural activity alone could account for the exquisite mapping that develops between body and brain. As is well known, peripheral manipulations of the visual, auditory, and somatosensory systems produce marked changes in cortical representations [21–24]. Nowhere is this more clear than in developing barrel cortex, in which whisker removal and peripherally expressed genetic manipulations reliably alter its structural organization (see [25] for review).

Nonetheless, recent reports appear to show that sensory feedback from movements is unrelated to ongoing neural activity within the somatosensory system [3, 4]. We suspected that these contradictory findings resulted from the use of methods that are inadequate for detecting movement-related neural activity. Thus, in the present study, we paid close attention to the physiological needs of the pups and ensured that they were cycling normally between sleep and wake and used methods with sufficient temporal resolution to ensure detection of movement-related neural activity. As a result, we provide here the strongest evidence to date that the majority of neural activity in barrel cortex is triggered specifically by whisker movements and that sleep is a critical modulator of this process.

By expanding our analysis to define periods when no movement was detected throughout the body, we found that the rate of spindle bursts was only 25 or 31% of that expected during active sleep or wake, respectively (Figure 5B). These values might be interpreted as an estimate of the “spontaneous” or “intrinsic” rate of spindle bursts, that is, spindle bursts that are independent of movement per se. Interestingly, these values are largely consistent with previous studies in which researchers blocked sensory feedback from the whiskers [11] or limbs [9, 12] and observed 50-75% decreases in the rate of spindle bursts. The current findings show that, by carefully attending to self-generated movement and behavioral state in intact animals, we can accurately assess the relationships between behavior and early brain activity.

The distinction between sensory input and “intrinsic activity” as contributors to cortical development was extended recently to the late prenatal period in mice (see [3]). Asserting that this is a period “before sensory input,” the authors concluded that sensory input cannot play a role in the observed cortical map structure. However, the circuitry that enables movement-related sensory feedback is already functional in thalamus by embryonic day 16 [26] and in cortex by embryonic day 18 [3]. Further, self-generated movements—as well as behavioral responses to cutaneous stimulation—are present by embryonic day 16 [27, 28]. Thus, it seems likely that sensory feedback drives subcortical neural activity long before the age tested here.

This study has several limitations. First, because we trimmed most of the contralateral whiskers and restrained the forelimbs, we were unable to determine whether movements of the trimmed whiskers and the forelimbs can account for any of the remaining residual activity. Second, we have found that the relationship between neural activity in developing barrel cortex and self-generated movements is probabilistic: Movements increase the likelihood of, but do not determine, a cortical response. In addition to behavioral state and movement amplitude, there are likely other contributing factors that remain to be identified. Finally, we found that cortical activity is greater during active sleep than wake, even during periods when pups were not moving (see Figure 5B). This finding suggests a role for state-dependent neuromodulation of corticothalamic excitability, a possibility that remains to be explored.

Altogether, our findings demonstrate that self-generated movements and sleep-wake state critically pattern early somatosensory cortical activity, including spindle bursts. In both visual and somatosensory cortex, the frequency, duration, and topographic specificity of spindle bursts are thought to be crucial in refining cortical topography through at least the first postnatal week [11, 13, 29]. Similarly, by triggering spindle bursts in somatosensory cortex, self-generated movements drive neural activity necessary for somatotopic development. Thus, future research should focus on the specific mechanisms through which patterned somatosensory input enables activity-dependent somatotopic development in this system.

## Acknowledgments

This research was supported by grants from the National Institutes of Health (R37-HD081168 to M.S.B. and F32-NS101858 to J.C.D.)

## Author Contributions

Conceptualization, J.C.D., R.M.G., and M.S.B.; Methodology, J.C.D., R.M.G., G.S., and M.S.B.; Software, J.C.D. and R.M.G.; Validation, J.C.D. and R.M.G.; Formal Analysis, J.C.D. and R.M.G.; Investigation, J.C.D. and R.M.G.; Resources, G.S. and M.S.B.; Data Curation., J.C.D. and R.M.G.; Writing - Original Draft., J.C.D. and R.M.G.; Writing - Reviewing and Editing, J.C.D., R.M.G., G.S., and M.S.B.; Visualization, J.C.D., R.M.G., G.S., and M.S.B.; Supervision, G.S. and M.S.B.; Project Administration, G.S. and M.S.B.; Funding Acquisition, G.S. and M.S.B.

## STAR METHODS

### CONTACT FOR REAGENT AND RESOURCE SHARING

Further information and requests for resources and reagents should be directed to, and will be fulfilled by, the lead contact, Dr. Mark Blumberg.

## SUBJECT DETAILS

Cortical activity was recorded in Sprague-Dawley Norway rats (*Rattus norvegicus*) at P5 (n = 8 pups from 8 litters; 5 males). In total, 37 single units were isolated (4.6 ± 0.64 single units per subject).

Animals were housed in standard laboratory cages (48 × 20 × 26 cm) on a 12:12 light-dark cycle, with food and water available ad libitum. The day of birth was considered P0 and litters were culled to eight pups by P3. All experiments were conducted in accordance with the National Institutes of Health (NIH) Guide for the Care and Use of Laboratory Animals (NIH Publication No. 80–23) and were approved by the Institutional Animal Care and Use Committee of the University of Iowa.

## METHOD DETAILS

### Surgery

As described previously [6, 30], a pup with a visible milk band was removed from the litter and anesthetized with isoflurane gas (3–5%; Phoenix Pharmaceuticals, Burlingame, CA). All whiskers on the left side of the snout, except those in the C row, were trimmed (Figure 1A). A custom-made bipolar hook electrode (0.002-inch diameter, epoxy coated; California Fine Wire, Grover Beach, CA) was inserted into the nuchal muscle for state determination. Carprofen (5 mg/kg SC; Putney, Portland, ME) was administered as an anti-inflammatory analgesic, and the animal’s forelimbs were wrapped in non-woven gauze to prevent them from stimulating the whiskers during the experiment.

After removing the skin above the skull, an analgesic was applied topically (bupivacaine; Pfizer, New York, NY). The skull was dried with bleach. Vetbond (3M, Minneapolis, MN) was applied to the skin around the skull and a head-plate (Neurotar, Helsinki, Finland) was secured to the skull using cyanoacrylate adhesive.

A trephine drill bit (1.8 mm; Fine Science Tools, Foster City, CA) was used to drill a hole into the skull above the right barrel field (0.5 mm posterior to bregma, 4.3 mm lateral from the sagittal suture). Two smaller holes were drilled distally to the recording site for insertion of a thermocouple and reference/ground electrode. A small amount of peanut oil was applied over the recording site to prevent drying of the cortical tissue. Surgical procedures lasted approximately 30 min.

The pup was then secured to a custom-made head-restraint apparatus inside a Faraday cage, with the animal’s torso supported on a narrow platform. Brain temperature was monitored using a fine-wire thermocouple (Omega Engineering, Stamford, CT) distal to the barrel field. The pup was allowed to recover from anesthesia and acclimate for 1 h. Recording did not begin until brain temperature was 36-37° C and the pup was cycling regularly between sleep and wake.

### Electrophysiological Recordings

The nuchal EMG electrode was connected to a Lab Rat LR-10 acquisition system (Tucker Davis Technologies, Gainesville, FL). The EMG signal was sampled at approximately 1.5 kHz and high-pass filtered at 300 Hz.

A 16-channel silicon depth electrode (Model A4×4-3mm-100-125-177-A16; NeuroNexus, Ann Arbor, MI) was coated in fluorescent Dil (Vybrant Dil Cell-Labeling Solution; Life Techologies, Grand Island, NY) before insertion. The electrode was inserted 500–800 μm into barrel cortex, angled 15-20° medially. A chlorinated Ag/Ag-Cl wire (0.25 mm diameter; Medwire, Mt. Vernon, NY) was inserted distal to the barrel field recording site, serving as both a reference and ground. The neural signal was sampled at approximately 25 kHz, with a high-pass (0.1 Hz) and a harmonic notch (60, 120, and 180 Hz) filter applied.

Electrode placement was confirmed using manual stimulation of the C-row whiskers. Neural activity from the barrel field was recorded for 1 h using SynapseLite (Tucker Davis Technologies) while the animal cycled between sleep and wake.

### Video Collection

In order to quantify whisker movements and identify behavioral states, we recorded video from two camera angles. Using two Blackfly-S cameras (FLIR Integrated Systems; Wilsonville, Oregon), the whiskers were filmed from below with the aid of an angled mirror and the body of the animal was filmed from the side. Video was collected in SpinView (FLIR Integrated Systems) at 100 frames per second, with a 3000 μs exposure time and 720×540 pixel resolution. The two cameras were hardwired to acquire frames synchronously and were initiated using a common software trigger.

The synchronization of video and electrophysiological data was ensured by using an external time-locking stimulus. A red LED controlled by SynapseLite (Tucker Davis Technologies) was set to pulse every 3 s for a duration of 20 ms. The LED was positioned to be in view of both cameras. Each video was analyzed frame-by-frame with custom Matlab scripts to ensure an equal number of frames between LED pulses. Although infrequent, when the number of frames between pulses was less than expected, the video was adjusted by duplicating and inserting one adjacent frame at that timepoint so as to preserve timing across the recording. Thus, all videos were ensured to be time-locked to the electrophysiological data within 10 ms.

### Histology

At the end of the recording period, the pup was euthanized with ketamine/xylazine (10:1; >0.08 mg/kg) and perfused with 0.1 M phosphate-buffered saline, followed by 4% paraformaldehyde. The brain was extracted and post-fixed in 4% paraformaldehyde for at least 24 h and was transferred to a 30% sucrose solution 24–48 h prior to sectioning. In order to confirm the electrode’s location within the barrel field, the right cortical hemisphere was dissected from the subcortical tissue and flattened between two glass slides (separated using two pennies) for 5-15 min. Small weights (10 g) applied light pressure to the top glass slide. The flattened cortex was sectioned tangentially to the pial surface. A freezing microtome (Leica Microsystems, Buffalo Grove, IL) was used to section the cortex (50 μm sections). Free-floating sections were imaged at 2.5x using a fluorescent microscope and digital camera (Leica Microsystems) to mark the location of the DiI.

Electrode placement in the C row of barrel cortex was confirmed by staining cortical sections for cytochrome oxidase (CO), which reliably delineates the divisions of primary sensory cortex at this age [31]. Briefly, cytochrome C (0.3 mg/mL; Sigma-Aldrich), catalase (0.2 mg/mL; Sigma-Aldrich) and 3,3’-diaminobenzidine tetrahydrochloride (DAB; 0.5 mg/mL; Spectrum, Henderson, NV) were dissolved in a 1:1 dilution of 0.1 M phosphate buffer and distilled water. Sections were developed in 24-well plates on a shaker (35–40°C, 100 rpm) for 3–6 h, rinsed in PBS, and mounted. The stained sections were imaged at 2.5x or 5x magnification and composited with the fluorescent image to confirm the electrode tract within the C row of S1 barrel cortex.

### Behavioral State and Whisker Movements

As described previously [5, 6], nuchal EMG and behavior were used to assess behavioral state (the experimenter was blind to the neurophysiological record when scoring behavior). The wake state was defined by the presence of high-amplitude movements of the limbs against a background of high nuchal muscle tone. Active sleep was defined by the presence of discrete myoclonic twitches of the hindlimbs and tail against a background of nuchal muscle atonia.

Whisker movements were quantified using DeepLabCut, a markerless tracking solution that uses a convolutional neural network to track features (e.g., whiskers) of animals in a laboratory setting [14, 15]. At least 200 manually labeled frames (tracking the tips of the whiskers) were used to initially train the network. After the initial training, newly analyzed frames with marker estimates that were deemed inaccurate were re-labeled and used to re-train the neural network until satisfactory tracking was achieved. Separate neural networks were trained for animals with three, four, and five visible whiskers (for best performance). The networks reached a training root mean square error (RMSE) of 0.03 mm for all networks and a test RMSE of 0.08, 0.09, and 0.07 mm, respectively.

Whisker twitches and wake movements were identified using custom Matlab scripts. Briefly, the y-axis (anterior-posterior axis) displacement of all C-row whiskers was averaged to obtain a single measure of whisker displacement. The onset of a whisker movement was determined as the first time point at which whisker displacement exceeded 3 standard deviations from baseline. Whisker movements that coincided with a jaw movement were initially separated from whisker movements that did not coincide with jaw movements. However, because no difference in the response magnitude of the neural activity was detected with or without jaw movements, all movements of the whiskers were included in the final analyses.

For measures of movement amplitude in Figures 3 and S3, whisker twitches and wake movements were normalized relative to maximum displacement for that subject. The amplitude of each whisker movement was then assigned to one of 10 equally sized bins.

To quantify movements of the ipsilateral whiskers, hindlimbs, and tail, we used custom Matlab scripts to detect frame-by-frame changes in pixel intensity within regions-of-interest (ROI). These frame-by-frame changes were smoothed using a moving Gaussian kernel with a half-width of 25 ms. Periods of movement for each ROI were determined by dichotomizing frame-by-frame changes with a threshold equal to the mean value across all frames; changes in pixel intensity above this threshold were defined as movement periods. When compared to experimentally scored movement periods, this threshold accurately separated periods of movement from periods of quiescence. Each period of movement began with the onset of the first movement, continued for as long as movements were no more than 500 ms apart, and ended 500 ms after the last movement.

### Local Field Potential

To assess local field potential power in relation to whisker movements, the neural signal was downsampled to 100 Hz and smoothed using a moving Gaussian kernel with a half-width of 0.5 ms. For time-frequency analyses, the LFP signal was convoluted using a Morlet wavelet (2–100 Hz using a linearly increasing range of cycles (2–10 cycles; [32]). Power is shown in dB, relative to the median power throughout a baseline period of 17 to 12 s before the movement. This baseline period was selected to achieve a separation between baseline and movement periods and to avoid edge effects within a ±20-s window.

To pinpoint spindle burst onset, the LFP was band-pass filtered at 8–40 Hz (stopband attenuation: -60 dB; transition gap: 1 Hz). The phase and amplitude of the filtered waveform were calculated using a Hilbert transformation. Spindle bursts were defined as an event for which the amplitude of the waveform was greater than the median plus 1.5 times the standard deviation of the overall waveform for 100 ms or longer. Once spindle bursts were isolated, the onset of each spindle burst was determined using a custom Matlab script.

### Spike Sorting

SynapseLite files were converted to binary files using custom Matlab scripts and sorted with Kilosort [33]. Briefly, the 16 channels of neural data were whitened (covariance-standardized) and band-pass filtered (300–5000 Hz) before spike detection. Next, template matching was implemented to sort the event waveforms into clusters. The first-pass spike detection threshold was set to 6 standard deviations below the mean and the second-pass threshold was set to 5 standard deviations below the mean. The minimum allowable firing rate was set to 0.01 Hz and the bin size for template detection set to 1,312,000 sample points, approximately 54 s. All other Kilosort parameters were left at their default values.

Clusters were visualized and sorted in Phy2 [34]. Putative single units (hereafter referred to as “single units” or “neurons”) were defined as having (1) spike waveforms that reliably fit within a well-isolated waveform template, (2) a distribution along seven principal components that was absent of clear outliers, and (3) an auto-correlogram with a decreased firing rate at time lag 0 (indicative of a single unit’s refractory period).

Clusters meeting the first two criteria but not the third were considered multi-units and were discarded from analysis. Any cluster with a waveform template indicative of electrical noise, a significantly low firing rate (< 0.01 spikes per second), or amplitude drift across the recording period was discarded.

## QUANTIFICATION AND STATISTICAL ANALYSIS

### Statistical Analysis

All data were tested for normality using the Shapiro-Wilk test and tested for sphericity with Mauchly’s test (where applicable). All data were normally distributed. Violations of sphericity were only observed for the main effect of amplitude, interaction, and simple main effects of the unit activity data presented in Figure S4, as well as the interaction in the data presented in Figure 5B (not significant). A Huynh-Feldt correction to the degrees of freedom for these analyses was performed.

Probabilities were arc-sin transformed to account for edge effects prior to analyses. A log_10_ transform was performed on the baseline rate of unit activity across state (Figure S1D).

All measures of central tendency and dispersion are represented as the mean and the standard error of the mean (SEM), respectively. In analyses of LFP and spindle burst activity, N refers to the number of animals (N = 8). In analyses of single-unit activity, N refers to the number of single units across animals (N = 37).

For direct comparisons between sleep and wake states, a univariate t-test was applied to the difference scores. For comparisons between pre- and post-movement rates of neural activity as well as behavioral state differences, a 2×2 repeated-measures ANOVA was performed. Throughout, interaction terms are reported, except in the analysis of Figure 4C (only the main effect of observed-expected likelihood was statistically significant) and Figure 5C (both the main effects of movement category and behavioral state were statistically significant). Simple main effects were always parsed by behavioral state, and a one-way ANOVA performed for both active sleep and wake.

All measures of effect size are reported as the adjusted partial-eta squared, an estimate of effect size that removes the positive bias due to sampling variability [35].

## DATA AND CODE AVAILABILITY

Custom MATLAB scripts used in this study can be found on Github (https://github.com/XXXXX). The data used to generate the figures in this paper are available upon request.

## Supplemental Information

**Figure S1.**
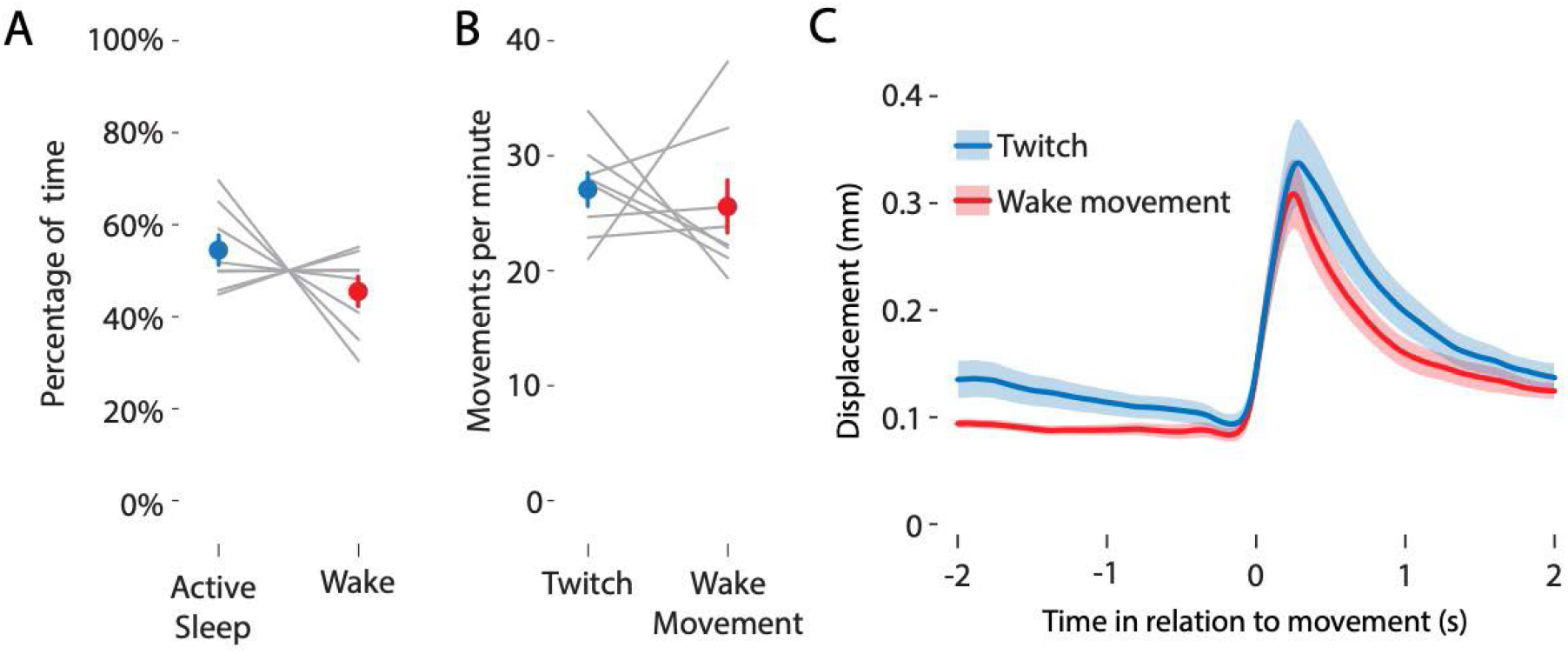
Behavioral state and movement data (A) Mean (±SEM) percentage of time spent in active sleep (blue circle) or wake (red circle). Gray lines show data for individual pups (N = 8). (B) Mean (±SEM) rate of whisker twitches (blue circle) and wake movements (red circle) per minute. Gray lines show data for individual pups (N = 8). (C) Mean (±SEM) displacement, in mm, for whisker twitches (blue line) and wake movements (red line) in relation to movement onset.

**Figure S2.**
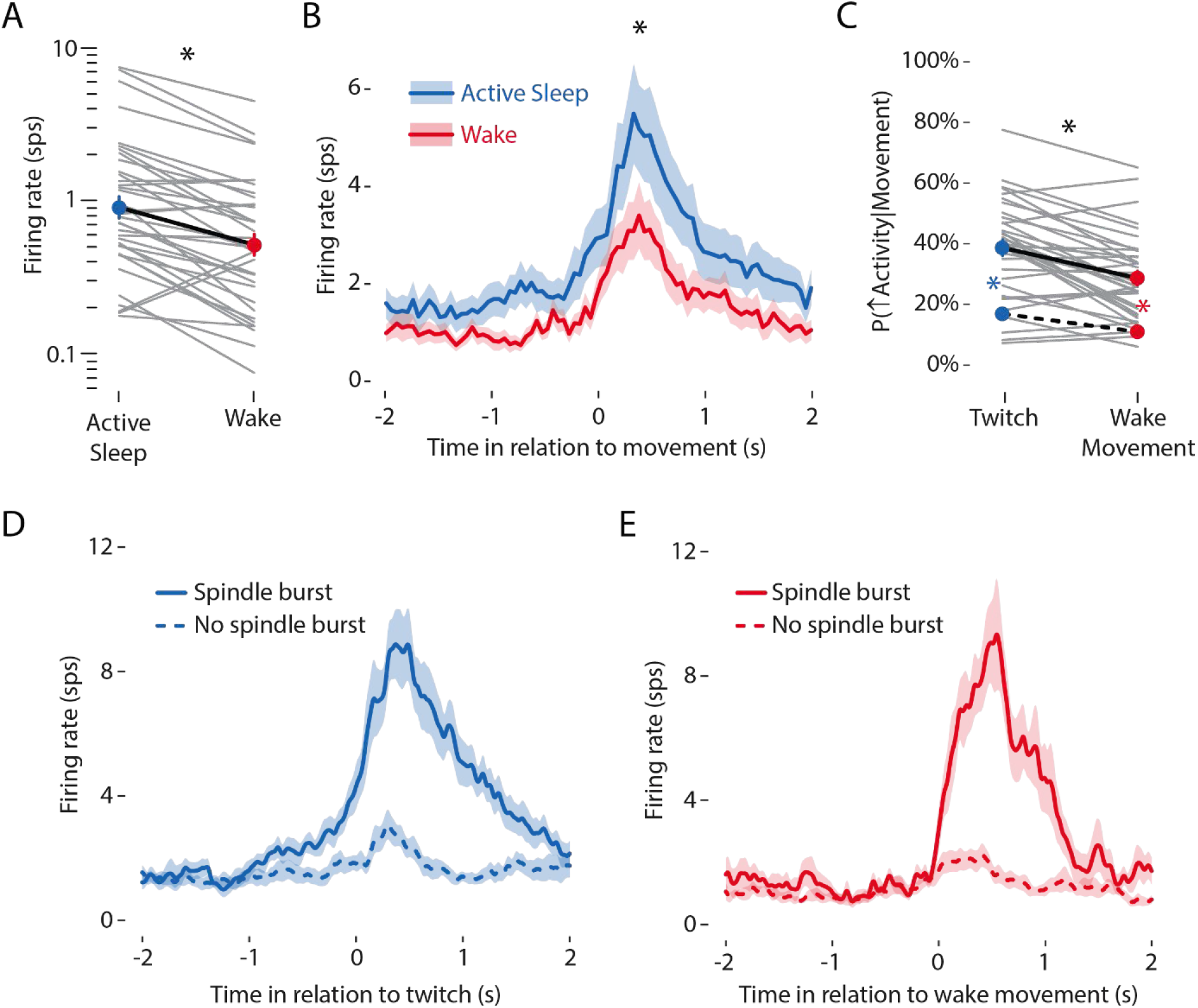
Whisker movements drive unit activity in barrel cortex (A) Mean (±SEM) firing rate of single units during active sleep (blue circle) and wake (red circle). Gray lines show the firing rate for individual units (N = 37). (B) Mean (±SEM) firing rate of all single units in relation to the onset of a twitch (blue line) or wake movement (red line). Asterisk indicates significant difference in peak firing rate between the two states (p < .05). (C) Mean (±SEM) likelihood of a firing rate increase given a whisker twitch (upper blue circle) or wake movement (upper red circle). The lower blue and red circles with dashed line indicate the mean (±SEM) expected likelihood due to chance during active sleep and wake, respectively. Gray lines show data for individual units (N = 37). The black asterisk denotes significant interaction (p < .05). Colored asterisks denote significant observed-expected likelihood differences (p < .05) for whisker twitches (blue) and wake movements (red). (D) Mean (±SEM) single-unit firing rate in relation to twitches that either did (solid line) or did not (dashed line) trigger a spindle burst. (E) Same as (D), but for wake movements.

**Figure S3.**
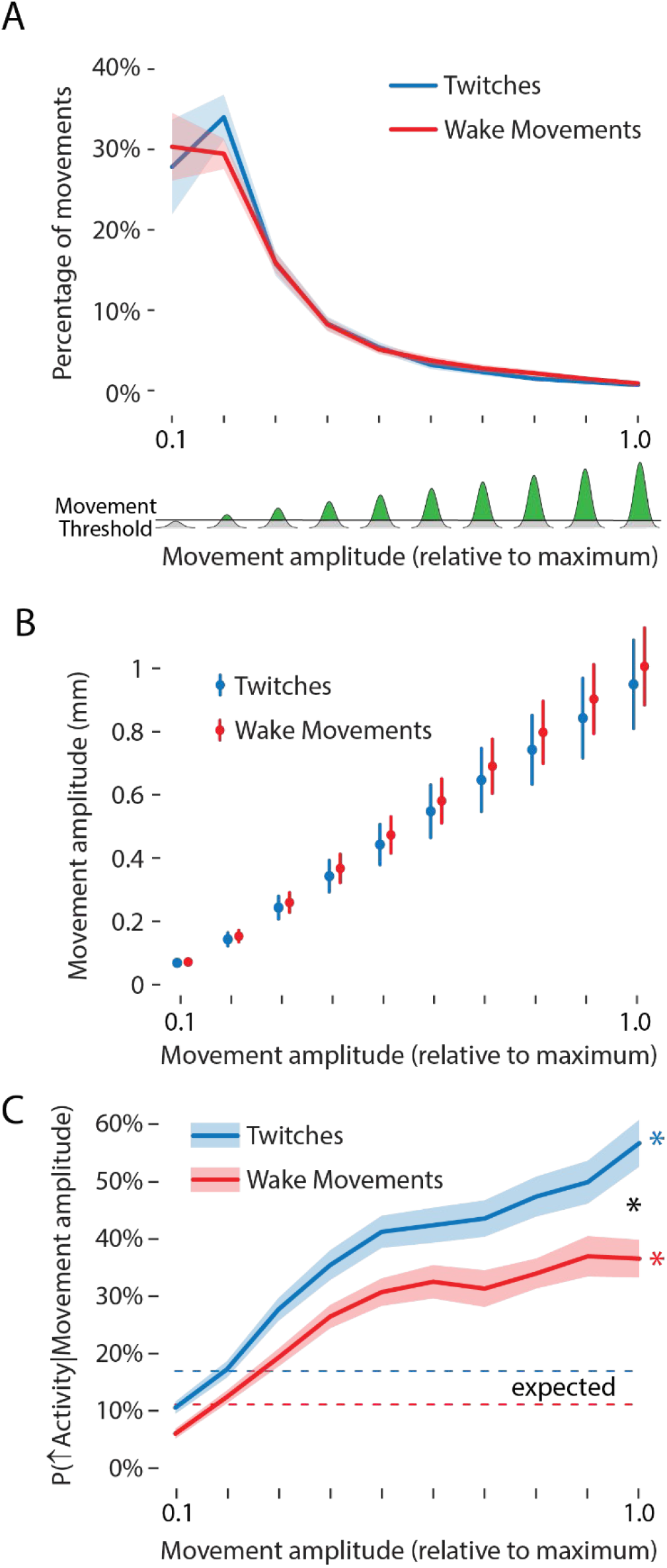
Movement amplitude moderates unit activity (A) Mean (±SEM) percentage of twitches (blue line) and wake movements (red line) sorted into 10 bins by movement amplitude, normalized to the maximum whisker amplitude for each pup. Amplitude is represented from 0 to 1 along the x-axis. Illustrated at bottom, in green, is the amount of whisker amplitude relative to the maximum for each pup; the horizontal line denotes the threshold used to detect whisker movements (see Methods). (B) Mean (±SEM) movement amplitude, in mm, of whisker twitches (blue circles) and wake movements (red circles) for each bin. The mean amplitude for each bin was not significantly different between twitches and wake movements. (C) Mean (±SEM) likelihood of a firing rate increase given a whisker twitch (blue line) or wake movement (red line) for each bin. The expected likelihood of a firing rate increase, for movements of any amplitude, is depicted as a dashed blue (active sleep) or dashed red (wake) line. The black asterisk denotes a significant interaction (p < .05). Colored asterisks indicate significant observed-expected likelihood differences (p < .05) for whisker twitches (blue) and wake movements (red).

**Figure S4.**
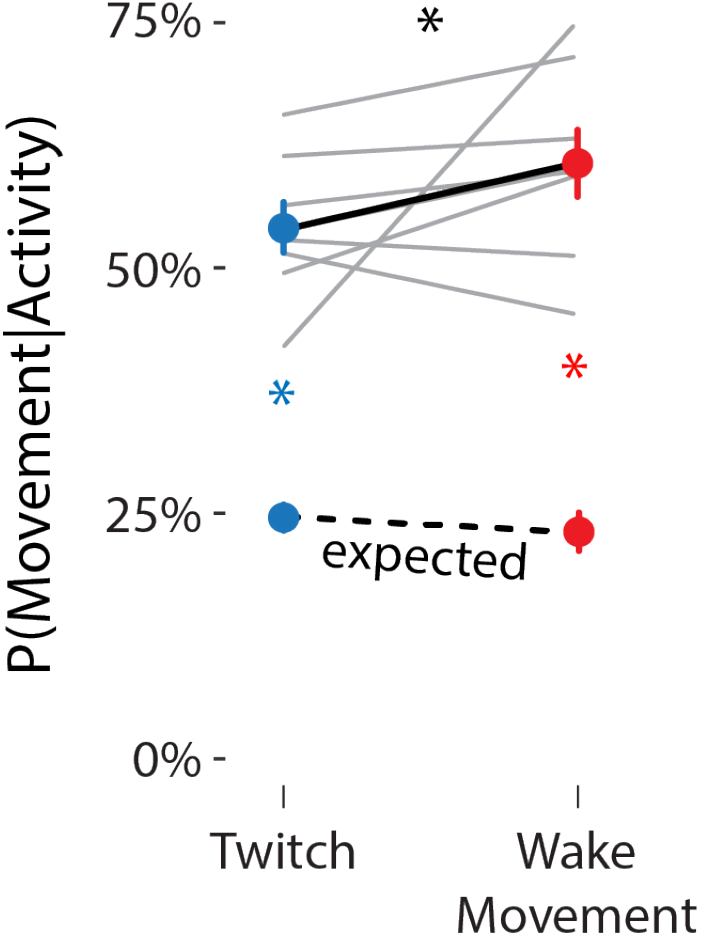
Percentage of unit activity attributable to self-generated movements Mean (±SEM) percentage of unit activity preceded within a 500-ms window by a whisker twitch (upper blue circle) or wake movement (upper red circle). The lower colored circles represent the mean (±SEM) likelihood of a twitch (blue) or wake movement (red) due to chance. Gray lines show data for individual pups (N = 8). The expected movement likelihood for individual pups is not shown. The black asterisk indicates a significant interaction (p < .05). Colored asterisks indicate significant observed-expected likelihood differences (p < .05) for whisker twitches (blue) and wake movements (red).

